# Winter temperature effects in a cold-adapted northern population of a range-expanding spider: survival, energy stores, and differential gene expression

**DOI:** 10.1101/2024.12.28.630119

**Authors:** Carolina Ortiz-Movliav, Marina Wolz, Michael Klockmann, Andreas Kuß, Lars Jensen, Corinna Jensen, Alexander Wacker, Gabriele Uhl

**Affiliations:** General and Systematic Zoology, Zoological Institute and Museum, University of Greifswald, 17487 Greifswald, Germany; Center for Functional Genomics of Microbes, University of Greifswald, 17489 Greifswald, Germany; Interfaculty Institute for Genetics and Functional Genomics, University of Greifswald, 17489 Greifswald, Germany; Animal Ecology, Zoological Institute and Museum, University of Greifswald, 17487 Greifswald, Germany

**Keywords:** cold-adaptation, range expansion, climate change, overwintering, energy stores, fatty acids, gene expression, RNA-seq, *Argiope bruennichi*, Araneae

## Abstract

Species expand their spatial distribution when environmental conditions are favorable or when mutations arise that allow them to live in previously unfavorable conditions. The European wasp spider, *Argiope bruennichi*, is known to have expanded its range poleward faster than climate change would predict. Northern edge populations show higher cold tolerance and are genetically differentiated from core populations, suggesting local adaptation to colder winter conditions. To investigate the degree and limits of plasticity in a cold-adapted population, we exposed overwintering juveniles (spiderlings) from Estonia - the northern edge of the distribution - to three winter regimes: two with a strong difference in day/night temperatures and an overall 10 degrees’ difference (warm and cold treatment) and one with moderate temperatures and less difference between day and night (moderate). We investigated if survival, lipid content, metabolites, and gene expression patterns differ depending on these temperature regimes. The survival probability of the spiderlings and their overall lipid content decreased over winter, with no difference between treatments, suggesting high resilience of the spiderlings towards very different temperature regimes at the edge of the distribution. At the end of winter, the content of saturated and monounsaturated fatty acids per spiderling also did not differ between treatments. However, omega-3 polyunsaturated fatty acids (PUFAs) levels were significantly lower in spiders exposed to the warm winter suggesting increased metabolic activity. We identified 4096 significant differentially expressed genes (DEGs) across the treatments, of which 1389 were specific for the moderate treatment, and 832 specific for the warm treatment, while 69 were unique for the cold treatment, showing a stronger temperature stress response to the moderate and warmer than to the cold treatment. Taken together, our results show that *A. bruennichi* has physiological plasticity and the ability to cope with very different winter temperature regimes despite being cold adapted. However, warmer winters might come with metabolic costs that could impact on the spiderlingś survival and foraging success when they emerge from the egg sac in spring.

## Introduction

With global climate change, temperatures have risen especially in the high-latitude regions, and will continue to do so under all climate change projections (Ono et al., 2022). Winter temperatures in these regions are also rising faster than in any other season (Hahn et al., 2023), leading to for example, reduced snow cover, potentially exposing overwintering arthropods to unexpectedly low winter temperatures (Bale & Hayward, 2010). Additionally, the increasing frequency of extreme temperature events (Renault et al., 2022) poses a significant threat to cold-adapted species (Shirey et al., 2024). Understanding how these temperature extremes affect the physiology of overwintering organisms will help assess and predict the resilience and survival in a changing climate.

Since the body temperature of spiders is strongly influenced by the ambient temperature (Harvey & Dong, 2023), and body temperature affects thermal performance patterns, it can be assumed that spiders are just as affected by climate change as other ectothermic animals (Burraco et al., 2020; Renault et al., 2022). However, since spiders have lower metabolic rates than expected from their body mass (Anderson, 1970; Schmitz, 2016), responses to temperature in spiders might differ from those of insects. For the latter a wealth of information is available (Bale & Hayward, 2010; Paaijmans et al., 2013; Harvey et al., 2020; Roberts et al., 2023). Increased knowledge on temperature susceptibility of spiders is needed to understand their thermal physiology and limitations in the face of climate change and for an informed comparison with insects.

The European Wasp Spider, *Argiope bruennichi* (Scopoli, 1772) (Araneae: Araneidae) lends itself for exploring how winter temperatures affect performance and physiology. This species has undergone a remarkably rapid poleward range expansion from the Mediterranean to Scandinavia and the Baltic region since the 1950s (Kumschick et al., 2011; Krehenwinkel et al., 2016; Wawer et al., 2017), a phenomenon commonly attributed to climate change (Hickling et al., 2006). However, the species has reached latitudes where its occurrence is difficult to explain by global warming alone (Geiser, 1997; Kumschick et al., 2011) as the speed of expansion outpaces the rate of warming. The newly colonized northern habitats have stronger seasonality, with shorter summers and colder winter conditions than the Mediterranean core area. Spiders from northern regions are already genetically differentiated from those of the core region (Krehenwinkel & Tautz, 2013, Sheffer et al. 2024, preprint). Adult spiders from the northern edge of the range having smaller body sizes and juveniles are less prone to disperse via ballooning (Wolz et al., 2020; Sheffer, 2024, preprint), juveniles prefer colder temperatures when given a choice (Krehenwinkel & Tautz, 2013) and can cope with significantly lower temperatures during winter as determined by lower lethal temperatures and lower super cooling points compared to juveniles from the core region (Sheffer et al. 2024, preprint). Thus, although the species original distribution is in the Mediterranean region, the populations in the north are already cold adapted and might therefore be vulnerable to warmer or moderate winter temperatures.

*A. bruennichi* overwinter as juveniles, so called spiderlings, in an intricately woven egg sac. They molt once before winter and remain in the shelter of the egg sac until they emerge from it in the spring of the following year, build orb-webs and continue their development until maturation in summer (Schaefer, 1977). Once the spiders live outside the egg sac, they can respond to ambient temperatures using behavioral measures, e.g. by changing the location of their webs. However, when inside the egg sac, the juveniles are subject to the winter conditions with no possibility of moving to a more benign microclimate. Therefore, the overwintering period represents the phase in which the vulnerability and physiological plasticity in response to temperature can be most accurately assessed (Williams et al., 2012).

During winter, physiological processes primarily focus on survival and preserving resources for growth in spring. This is especially the case in animals that are unable to replenish energy stores during this period and are thus limited to a fixed energy budget. Storage lipids (triacylglycerol fatty acids) serve as the primary energy source during winter (Hahn & Denlinger, 2011; Sinclair & Marshall, 2018). In cold-adapted overwintering species, specific modifications in lipid compositions are observed, including a higher ratio of unsaturated to saturated fatty acids (Storey & Storey, 2013). This pattern is also evident in the fatty acids incorporated into cell membrane phospholipids. By increasing the proportion of mono- and polyunsaturated fatty acids in the membrane phospholipids, the membrane homeoviscosity can be maintained in colder environments (van Dooremalen & Ellers, 2010, Sinclair & Marshall, 2018).

Low temperatures have led to the evolution of specific cold-tolerance mechanisms (Storey & Storey, 2013; Sinclair et al., 2015). Freeze avoidant species, such as *A. bruennichi*, produce cryoprotectants and antifreeze proteins that shift the point of freezing to lower temperatures (mean -28.75°C, in spiderlings from the edge that overwinter in cold conditions) (Sheffer, 2024, preprint). The substances involved in deep supercooling are considered costly to produce (Storey & Storey, 2013) and their accumulation can also increase hemolymph viscosity (Vanin et al., 2008), thereby negatively impacting lipid transport (Sinclair & Marshall, 2018). On the other hand, in warm phases during the winter period the metabolic rate goes up and energy stores are quickly diminished (Clarke & Fraser, 2004; Sinclair 2015; Roberts et al. 2023). Especially in ectotherms, even small changes in ambient temperature can lead to considerable physiological costs and biochemical adjustments (van Dooremalen & Ellers, 2010).

Our study aims at investigating if a recently cold-adapted population of *A. bruennichi* at the edge of the range is resilient or susceptible to winter regimes that differ in overall temperature and in the degree of daily temperature fluctuation. We exposed spiderlings from Estonia to cold (5°C day, -15°C night), moderate (5°C day, -5°C night), and warm (15°C day, -5°C night) winter temperature regimes over three months. We measured how the spiderlingś survival and physiology is affected by the temperature regime over time by opening egg sacs at two time points (after 1 and 3 months). We assessed survival rate and the total lipid content to gain insight into the dynamics of the spiderlingś performance and usage of energy stores under strongly different temperature regimes. After three months’ exposure, we also explored the fatty acids profiles and investigated if specific gene families are differentially expressed depending on the temperature treatment. We predicted that temperature regimes would impact the spiderlingś performance and physiological traits and that this would be reflected in differential gene expression patterns.

## Material and Methods

### Sampling and winter treatments

Adult mated female spiders of *Argiope bruennichi* (NCBI: txid94029) were collected in August 2017 from the edge of the distribution in Pärnu, Estonia (N: 58.294, E: 24.602). After collection, we brought the spiders into a climate-controlled chamber at the University of Greifswald, Germany, and kept them under shared conditions (20°C, 70% relative humidity) until oviposition. The females were housed in individual plastic containers (1000 mL) (n = 170) where they laid eggs into a silk-woven egg sac (Fig. 1A). The females were misted with water daily and fed regularly with house flies (*Calliphora* sp.). After a second oviposition, females were anesthetized with CO_2_ and preserved in ethanol. The egg sacs remained in the climate chamber in individual 5 x 5 x 3.5 cm plastic containers with mesh on opposite sides for ventilation. Inside the egg sacs, the post-embryos hatch from the eggs already a few weeks after oviposition and the post-embryos molt once to first instar spiders (spiderlings) that remain in the egg sac over winter (Leborgne & Pasquet, 2005). The spiderlings emerge from the egg sac in spring of the following year and start building webs near the ground.

**Figure 1.**
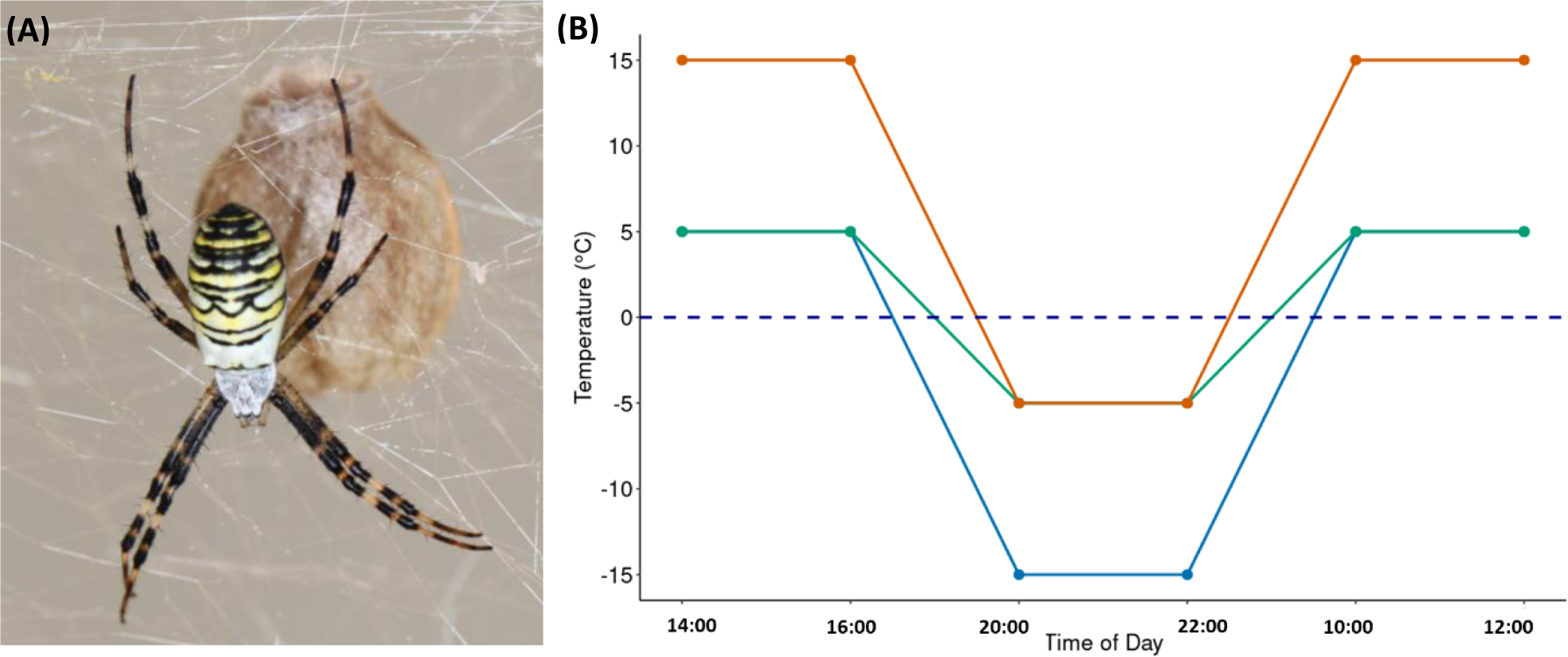
**A.** *Argiope bruennichi* female with a silk woven egg sac, photo by G. Uhl **B.** Daily temperature simulations in a one-day cycle. Blue line = cold treatment (5 °C day/-15 °C night; n = 49), green line = moderate treatment (5 °C day/ -5 °C night) and orange line = warm treatment (15 °C day/ -5 °C night; n = 52).

**Figure 1.**
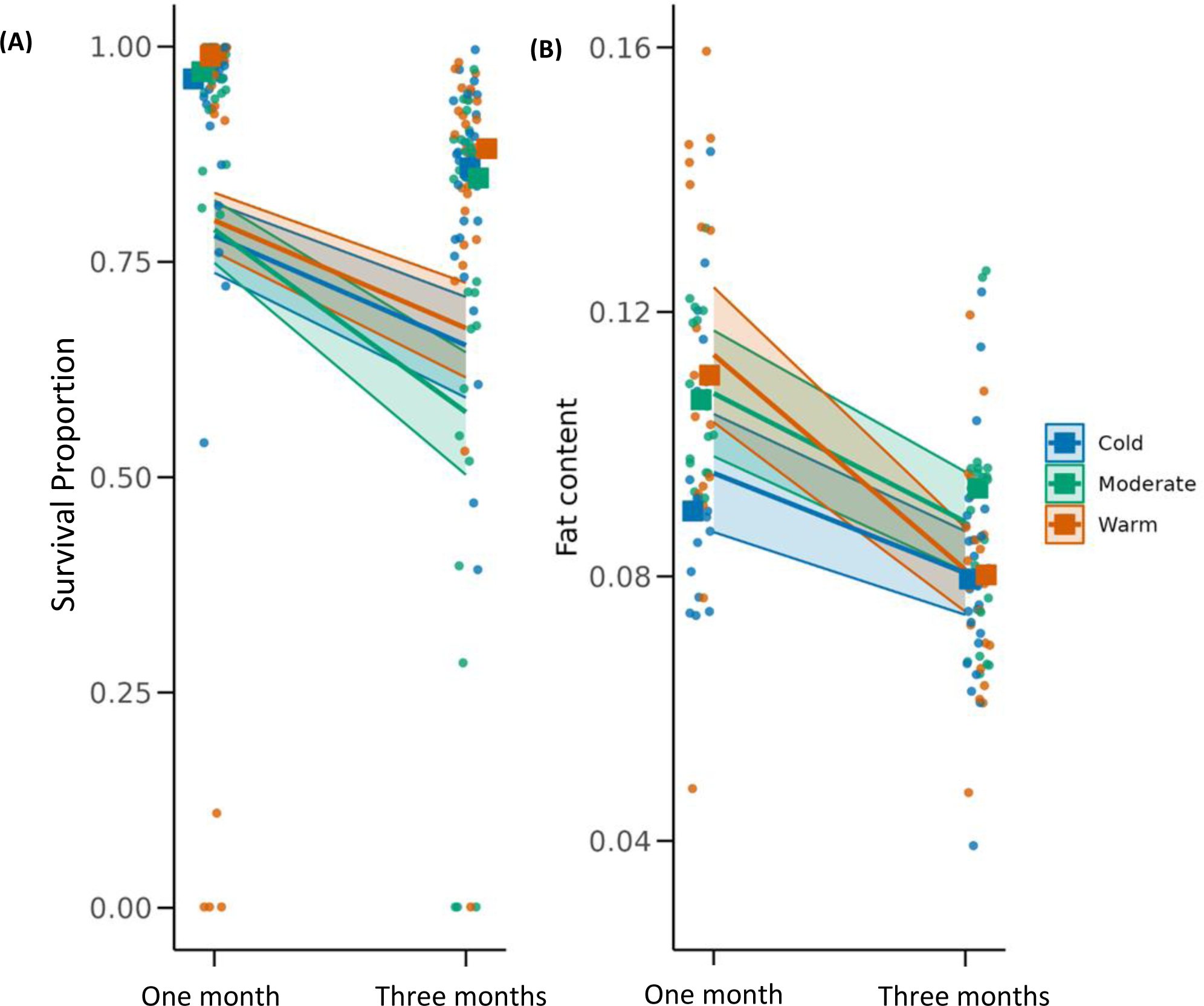
**Temperature related differences in *Argiope bruennichi* spiderlings** in the **(A)** Proportion of live offspring (survival) after one month and three months of exposure to winter treatments **(B)** Fat content of spiderlings after one month and three months of exposure to winter treatments. Lines represent model predictions flanked with their 95% confidence intervals, the points represent the raw data, and the squares represent the median of the data.

On November 17^th^ 2017, 153 egg sacs with spiderlings inside were randomly distributed to three climate cabinets simulating a cold winter (5 °C day/-15 °C night; n = 49; Percival LT-36VL, CLF PlantClimatics GmbH, Wertingen, Germany), moderate winter (5 °C day/ -5 °C night; n = 52), and warm winter (15 °C day/ -5 °C night; n = 52; Fig.1). Eggs sacs of warm and moderate winter regimes were kept in Panasonic MLR-352H climate cabinets (Ewald Innovationstechnik GmbH, Bad Nenndorf, Germany).

The egg sacs containing spiderlings were sprayed each evening with water to simulate increasing humidity at night. To assess how temperature affects survival during winter, we opened egg sacs at two time points, a month after the start of the winter treatment (December 21^st^, 2017, cold n=21, moderate n =24 and warm n=34), and three months after the start of the winter treatments (February 19 - 23rd, 2018, cold n = 28, moderate n=28 and warm n=28). The number of live spiderlings, dead spiderlings as well as eggs were counted. We determined the survival proportion of the offspring as the number of live spiderlings divided by the total number of hatched spiderlings (live plus dead). The live spiderlings were flash-frozen at -80 °C and stored until further processing. Fat content was assessed for both exposure times whereas fatty acids and differential gene expression were analyzed after 3 months’ exposure.

### Fat content

From a subset of the total egg sacs (n= 125 out of 153), 30 spiderlings per egg sac were used for fat content analysis (after a month: n= 17 cold, n = 18 moderate, n = 17 warm; after three months: n= 26 cold, n = 21 moderate, n = 26 warm). The fat content was obtained following the protocol described in Geiger et al. (2018). The samples were dried for 48 h at 60°C and then weighed (initial dry weight) using a high-precision balance (Sartorius ME5, Sartorius AG, Gottingen, Germany). Subsequently, fat was extracted using 1 mL of acetone: After 48 h, the acetone was changed and left for an additional two days before removal. Then, the samples were again dried at 60°C for 48 h and subsequently weighed. Fat content was calculated as the mass difference between the two measurements. Relative fat content was calculated by dividing fat content by the initial dry weight.

### Fatty acid analysis

Lipids and fatty acids extraction was done for spiderlings from egg sacs that had experienced 3 months of exposure to one of the three temperature regimes. Eight samples were randomly chosen from each treatment, each containing 15 spiderlings. The lipids were first extracted by adding the samples to a glass vial with 4 mL of dichloromethane-methanol (2:1 v/v) and 50 µL nonadecanoic acid methyl ester (C19:0 200 ng/μL) as an internal standard (Wacker et al., 2016). After storage for 48 hours at -20 °C, samples were placed in an ultrasonic bath for 7 seconds, and the solvent was transferred to a new vial. A fresh dichloromethane-methanol (2:1 v/v) solution was added, and the closed vial placed into the ultrasonic bath for 20 seconds. Next, the solvent was collected and added to the solvent from the first extraction. The solvent was removed under a gentle stream of nitrogen gas by placing the vials in a concentrator evaporator thermostatically controlled at 40°C (Cole-Parmer™ Stuart™ SBH130D/3) to prevent lipid reaction to atmospheric oxygen. The dried samples were resuspended in 4 mL 3mol/L methanolic HCL (Sigma-Aldrich Chemie, Darmstadt, Germany) and incubated for 20 min at 60°C in a closed vial to transesterify fatty acids into fatty acid methyl esters (FAMEs). After cooling at 4°C for 20 min, FAMEs were extracted three times. Each time, 1 mL of isohexane was added to the vials containing the samples in methanolic HCl and vortexed for 5 seconds before allowing the two phases to settle. This vortexing step was repeated three times. The isohexane phases of the three extractions were collected in a new vial. Subsequently, this fraction was evaporated to dryness under nitrogen at 40°C, resuspended in 100 μL of isohexane, transferred to GC vials with micro inserts, and stored at -20°C until injection. The FAMEs were analyzed by gas chromatography (6890N, Agilent Technologies, Böblingen, Germany) according to Wacker & Weithoff (2009). Helium was used as carrier gas (1.5mL/min), and FAMEs were separated on a DB-225 column (Agilent, 30 m x 250 μm x 0.25 μm), applying an initial oven temperature of 60°C for 1 minute, increasing at a rate of 20°C/min until 150°C, 10°C/min until 220°C and then held for 14 min. FAMEs were detected using a flame ionization detector (FID) at 250°C and quantified using multipoint standard calibration curves with HP chemstation (Agilent Technologies). To check the identity of the fatty acid methyl esters, mass spectra were recorded with a gas chromatograph (7890A, Agilent Technologies) connected to a mass spectrometer (Pegasus 4D GC-TOFMS, LECO Instruments, Mönchengladbach, Germany). Data handling was carried out using associated chemstation software (6890N, Agilent technologies) and as described in Wacker et al. (2016). Fatty acids were identified on the basis of their retention time and compared to a Supelco standard (Supelco 37 Component FAME Mix) and quantified on the basis of multipoint calibration curves. To obtain the respective fatty acids contents per individual spiderling, we divided the total amounts by 15, i.e., the number of spiderlings used in each sample.

### Statistical analyses

The statistical analyses were performed with R version 4.3.1 (R core Team, 2023). In order to assess differences in survival and fat content of the spiderlings depending on the winter treatment and length of exposure, our explanatory variables were “winter treatment” (cold, moderate or warm), and the two time points of inspection, “exposure time”. We added the females’ ID as a random effect, to account for the use of second egg sacs (48 %) from one female, specified as (1 | Mother ID). We analyzed the survival proportion (logit transformed) and fat content using generalized linear mixed models, with the package glmmTMB (Brooks et al., 2017). We used models that assumed a tweedie family function, suitable for continuous response variables with zero inflation and over dispersion. Model assumptions were checked using the package DHARMa (Hartig & Lohse, 2022), and we used a type III ANOVA of the ‘car’ package and a post-hoc test with Tukey correction to test for the significance of the treatment effects on the response variables with the package emmeans (Lenth et al., 2024). To analyze differences between means of the different fatty acid groups an analysis of variance (ANOVA) was applied when the data were normally distributed, followed by a post-hoc Tukey test.

### RNA isolation and sequencing

For each winter treatment we used four replicates each consisting of 15 spiderlings (total n = 12). The pool of spiderlings in each sample were mechanically homogenized using non-sticky pellet pestles in the presence of 500 μL RNA-Solv Reagent (Omega Bio-tek, Norcross, GA, USA), following the manufacturer’s recommendations. The isolated RNA was dissolved in 20 μL of Nuclease-free water and quantified using a Qubit 2.0 (Life Technologies). RNA integrity was visualized with a Bioanalyzer 2100 system (Agilent, Santa Clara, CA, USA). 2 μg total RNA from each sample was subject to ribosomal depletion (rRNA) using a custom designed *Argiope bruennichi* riboPOOL depletion kit dp-K096-000074 (siTOOLs Biotech, Germany) following the manufacturer’s indications. The ribosomal depleted RNA was purified using the ethanol precipitation option and quantified using the Bioanalyzer 2100 system. We took 100 ng of ribosomal depleted RNA for library construction using the NEBNext® Ultra™ II Directional RNA Library Prep Kit for Illumina® (New England Biolabs, Ipswich, MA, USA). Sequencing was done on a NextSeq® 500/550 System using a High Output Kit v2.5 flow cell (150 cycles) and 2 x 75 bp paired-end sequencing.

### Differential gene expression analysis

We used fastqc 0.11.9 (http://www.bioinformatics.babraham. ac.uk/projects/fastqc) to check the quality of the raw data. After quality control, we mapped the reads to the genome of the species (accession No. GCA_947563725.1) (Sheffer et al., 2021) using the program STAR 2.7.11a (Dobin et al., 2013) in two-pass mode with default parameters. We used the resultant BAM files to generate a count table on Subread 2.0.6, using the feautureCounts function (Liao et al., 2014). The count data was normalized with the R package DESq2 1.38.3 (Love et al., 2014) and analyzed via principal components using the plotPCA function. We then ran three tests with these normalized data: (1) cold treatment vs moderate treatment, (2) moderate treatment vs warm treatment, and (3) cold vs warm treatment. Transcripts were differentially expressed if the Benjamini and Hochberg adjusted false discovery rate (FDR) p-value was less than 0.05. Overlaps of differentially expressed genes and unique gene sets across the winter treatments were calculated and visualized using a VennDiagram package (H. Chen, 2022). To determine whether the overlaps between different treatment pairs (cold vs. moderate, moderate vs. warm and cold vs. warm) were greater than would be expected by chance alone, Fisher’s Exact Tests were performed. Gene ontology (GO) enrichment analysis was performed with the package topGO version 2.56.0 (Rahnenfuhrer, 2024) to infer the biological function of all differentially expressed genes at a FDR of 0.05. Additionally, a Kyoto Encyclopedia of Genes and Genomes (KEGG) pathway enrichment analysis (corrected p < 0.05) was tested with clusterProfiler version 4.10.0 (Wu et al., 2021) using the *Argiope bruennichi* genome sequence and assembly (Assembly: GCF_947563725.1 chromosome)(Crowley et al., 2023). The most enriched, differentially expressed pathways, which can be divided into upregulated and down-regulated genes, were visualized using a heat map created with the ComplexHeatmap package (Gu, 2022).

## Results

### Survival and fat content

The proportion of spiderlings found alive was significantly higher after one month of temperature exposure (89 %) compared to 3 months (78 %) over all treatment groups (GLMM, χ^2^ = 12.61, p = 0.004, Fig. 2A). There was no significant difference in survival depending on winter treatment (GLMM, χ^2^ = 0.45, p = 0.80, Fig. 2A) The interaction between Treatment and Exposure Time was not significant (GLMM, χ^2^ = 2.62, P = 0.27).

**Figure 2.**
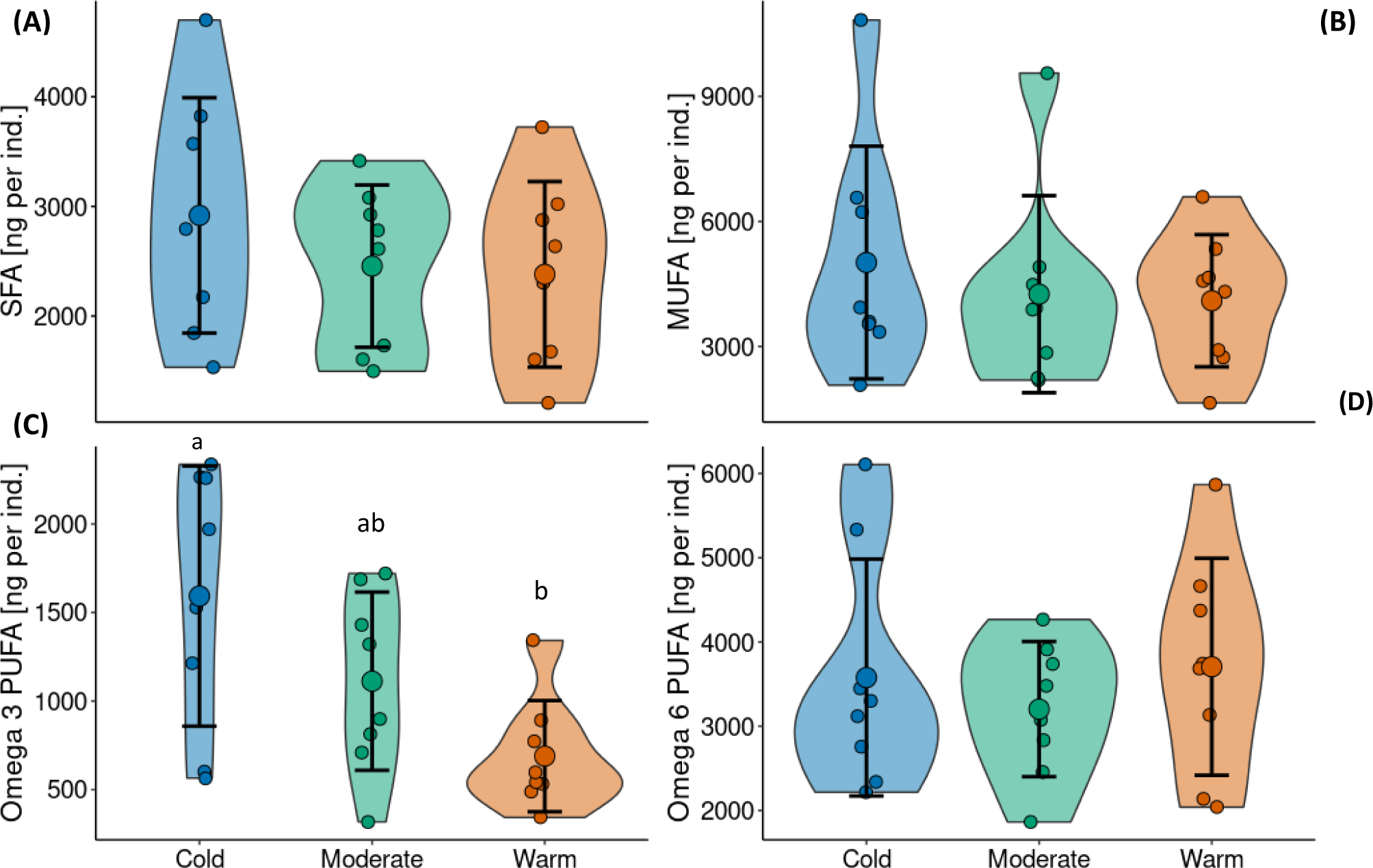
**Amounts of main fatty acids groups on a per carbon basis (ng fatty acid per individual)** of spiderlings according to winter treatment (cold, moderate, warm) in **(A)** Saturated fatty acids (SFA), **(B)** Monounsaturated fatty acids (MUFA), **(C)** Omega-3 polyunsaturated fatty acids (PUFA) and, **(D)** Omega-6 polyunsaturated fatty acids (PUFA). Large circles with error bars represent means ± s.d. Small circles represent the data points and the violins represent the distribution of the data. Different lowercase letters indicate significant differences among treatments (Tukey HSD post-hoc test following ANOVA).

Following the same pattern, the fat content of the spiderlings was significantly lower in the 3-month (8.3 %) compared to 1-month exposure group (10.6%) over all treatment groups (GLMM, χ^2^ = 8.48, p = 0.003, Fig. 2B). There was no significant difference in fat content depending on winter treatment (GLMM, χ^2^ = 4.13, p = 0.12, Fig. 2B). The interaction between Treatment and Exposure Time was not significant (GLMM, χ^2^_2_ = 2.33, P = 0.31).

### Fatty acids

The amount of saturated fatty acids, MUFA and omega-6-PUFAs in the spiderlings from the different overwintering treatments were not significantly different after 3 months of exposure (ANOVA, p > 0.05; Tab. S1). Also, the ratio between unsaturated and saturated fatty acids did not differ between treatments (ANOVA, F_2,21_= 0.21, p= 0.81; Tab. S1). Likewise, the amount of eicosapentaenoic acid (EPA, 20:5n-3) was largely constant within the range from 352 - 368 ng across all winter treatments (ANOVA F_2,21_=0.03, p = 0.97; Tab. S1). However, the spiderlings of the warm winter treatment contained 57% less omega-3 PUFAs compared to those from the cold winter treatment (Fig. 2C, Tab. S1). Spiderlings from the moderate treatment showed intermediate amounts. This difference between the warm and cold treatment was mainly caused by a 5.4 lower amount of α-linolenic acid (ALA, C18:3n-3) in the warm compared to the cold winter treatment (ANOVA F_2,21_= 6.7, p < 0.01; Tab. S1).

### Differential gene expression depending on winter temperature

We screened for genome-wide differential gene expression based on RNA-Seq data. From a total of 37,459 transcripts, we identified 14,893 differentially expressed genes (DEGs) between the three winter treatments after filtering for low TPM (Transcripts Per Kilobase Million) values and collapsing duplicates. Gene expression patterns differed between treatments, with PC1 accounting for 49% of the variation (Fig. 3A, Fig. S1). Expression patterns were distinctly different between the cold and warm treatments and highly variable in the moderate treatment. We identified 4,096 differentially expressed genes (DEGs) with a strong increase in expression (≥10 fold-change). Among the DEGs, 2,251 genes were upregulated and 1,845 downregulated. The warm winter treatment showed a higher overall transcriptomic response compared to the cold treatment, as demonstrated by significant effects in cold versus warm and moderate versus warm comparisons (Fisher’s exact test, two-tailed P < 0.001) (Table S2). Specifically, 832 DEGs were unique to the warm winter treatment, while only 69 were unique to the cold treatment. The moderate treatment had 1,389 unique DEGs, and 8 DEGs were common to all treatments (Fig. 3B). The highest contrast in the number of up- and downregulated genes was found in the cold vs. warm comparison, with 47 highly significant downregulated and 22 upregulated genes (Fig. S2C).

**Figure 3.**
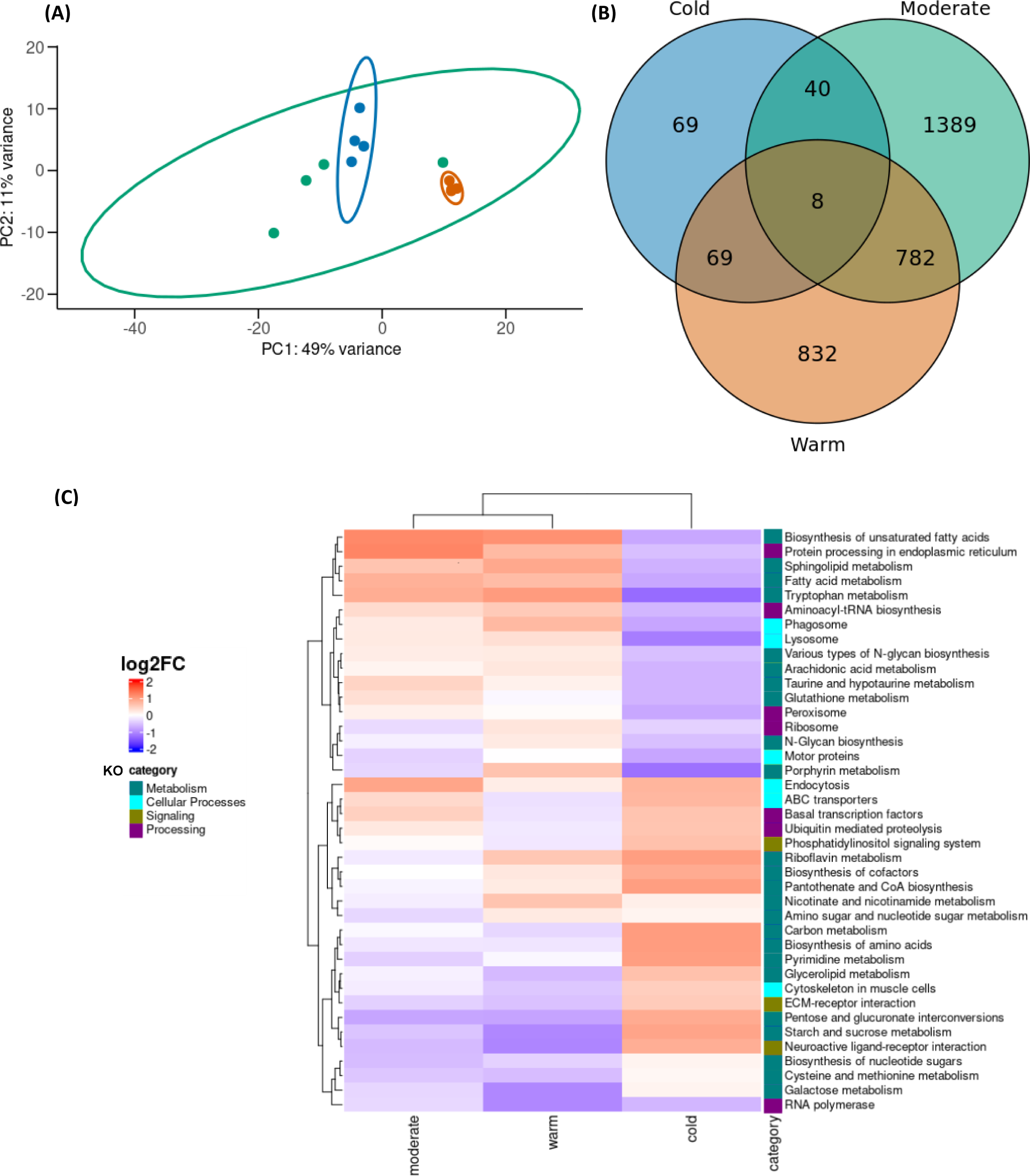
**Transcriptomic differences between the winter treatments. (A)** Principal component analysis of all differentially expressed transcripts (DETs) (n= 14893), on normalized data counts from 12 RNA-seq samples constituted of 15 spiderlings pooled for the three winter treatments: cold (blue), moderate (green) and warm (orange). The points represent the data, and the ellipses represent the confidence level of 95%. **(B)** Numbers of differentially expressed transcripts (FDR > 0.05) for treatment specific expression, as well as overlap between treatments. **(C)** Gene expression profile. Clustered heatmap of the top 40 enriched KEGG Orthology (KO) classes in *A. bruennichi* (abru), showing up and down regulated pathways between the treatments. Red indicates upregulation and blue downregulation based on the log2 fold-change.

The Gene Ontology (GO) term enrichment, similar to the gene expression pattern, was treatment-specific and had less enrichment in the cold treatment (Table 1). GO terms were identified in the categories Biological Processes (BP), Molecular Function (MF), and Cellular Component (CC), involved a total of 314 terms (Table 1 shows the 35 most enriched, for discussion within the text). The moderate treatment exhibited the highest degree of functional enrichment, followed by the warm and cold treatments. In the warm treatment, upregulated GO terms are related to calcium channel activity (MF), establishment of localization (BP), and transport (BP), all processes that are known to be involved in heat stress responses. In the cold treatment, downregulated terms included lipid modification and glutathione catabolic process (BP) (Table 1). GO terms enriched in the cold treatment (Table 1) are related to structural constituents of cytoskeleton (MF), cell communication, and signaling. The moderate treatment displayed enrichment for a large number of terms, including those associated with stress responses in general, such as phosphorus metabolic processes, homeostatic processes, and negative regulation of biological processes (BP) (Table 1).

**Table 1.**
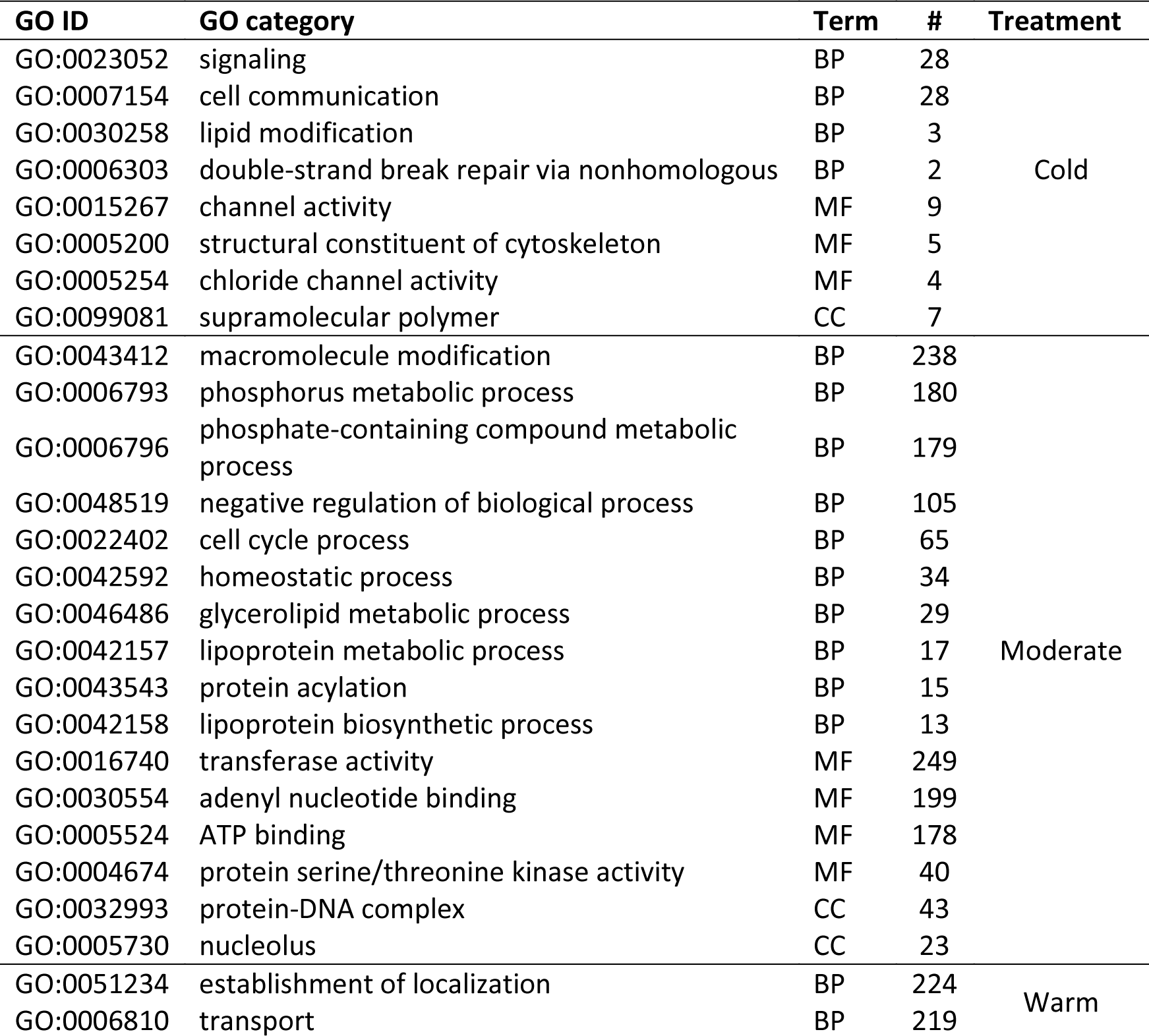
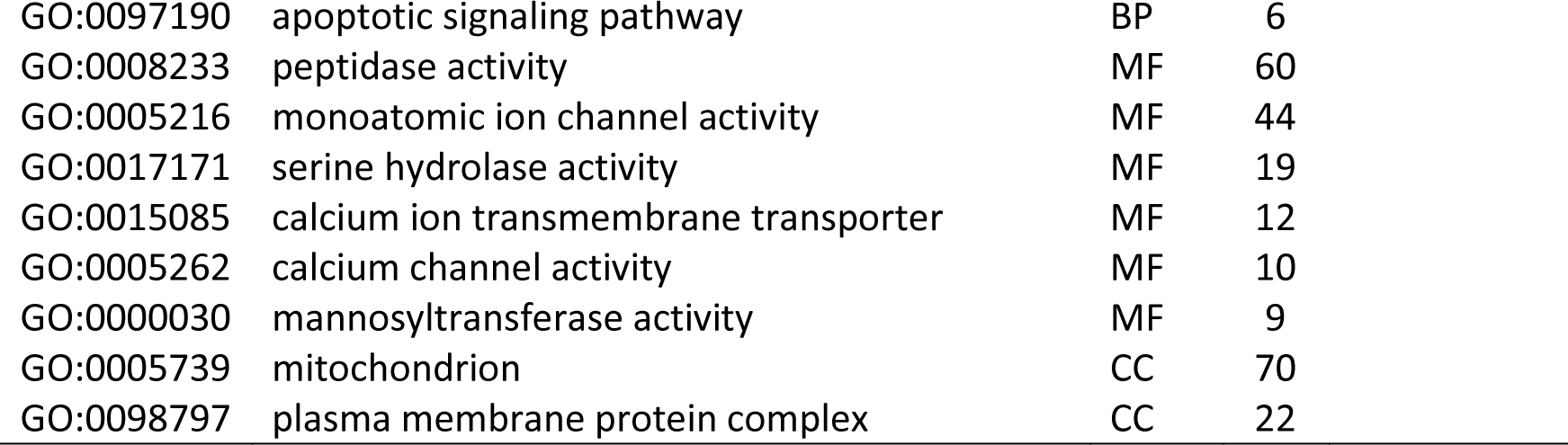
Significantly enriched Gene Ontology (GO) categories for differentially expressed genes in the cold, moderate or warm winter treatments at a false detection rate (FDR) of 0.05. # refers to the number of enriched transcripts in the category. BP = Biological processes, MF= Molecular function and CC = cellular component.

| GO ID | GO category | Term | # | Treatment |
| --- | --- | --- | --- | --- |
| GO:0023052 | signaling | BP | 28 | Cold |
| GO:0007154 | cell communication | BP | 28 |  |
| GO:0030258 | lipid modification | BP | 3 |  |
| GO:0006303 | double-strand break repair via nonhomologous | BP | 2 |  |
| GO:0015267 | channel activity | MF | 9 |  |
| GO:0005200 | structural constituent of cytoskeleton | MF | 5 |  |
| GO:0005254 | chloride channel activity | MF | 4 |  |
| GO:0099081 | supramolecular polymer | CC | 7 |  |
| GO:0043412 | macromolecule modification | BP | 238 | Moderate |
| GO:0006793 | phosphorus metabolic process | BP | 180 |  |
| GO:0006796 | phosphate-containing compound metabolic process | BP | 179 |  |
| GO:0048519 | negative regulation of biological process | BP | 105 |  |
| GO:0022402 | cell cycle process | BP | 65 |  |
| GO:0042592 | homeostatic process | BP | 34 |  |
| GO:0046486 | glycerolipid metabolic process | BP | 29 |  |
| GO:0042157 | lipoprotein metabolic process | BP | 17 |  |
| GO:0043543 | protein acylation | BP | 15 |  |
| GO:0042158 | lipoprotein biosynthetic process | BP | 13 |  |
| GO:0016740 | transferase activity | MF | 249 |  |
| GO:0030554 | adenyl nucleotide binding | MF | 199 |  |
| GO:0005524 | ATP binding | MF | 178 |  |
| GO:0004674 | protein serine/threonine kinase activity | MF | 40 |  |
| GO:0032993 | protein-DNA complex | CC | 43 |  |
| GO:0005730 | nucleolus | CC | 23 |  |
| GO:0051234 | establishment of localization | BP | 224 | Warm |
| GO:0006810 | transport | BP | 219 |  |

|  |  |  |  |
| --- | --- | --- | --- |
| GO:0097190 | apoptotic signaling pathway | BP | 6 |
| GO:0008233 | peptidase activity | MF | 60 |
| GO:0005216 | monoatomic ion channel activity | MF | 44 |
| GO:0017171 | serine hydrolase activity | MF | 19 |
| GO:0015085 | calcium ion transmembrane transporter | MF | 12 |
| GO:0005262 | calcium channel activity | MF | 10 |
| GO:0000030 | mannosyltransferase activity | MF | 9 |
| GO:0005739 | mitochondrion | CC | 70 |
| GO:0098797 | plasma membrane protein complex | CC | 22 |

KEGG enrichment analysis revealed that the DEGs could be classified into 121 pathways (Table 2 shows the 36 most enriched). The top 40 highly enriched up and downregulated terms are shown in Figure 4C. The KEGG pathways were identified in four categories or KO classes (KEGG ontology classes): Metabolism, Cellular processes, Signaling and Processing. The Metabolism were the most enriched pathways overall. The most enriched metabolism pathways included those involved in glycine, serine, and threonine metabolism; carbon metabolism; glycerophospholipid metabolism; fatty acid metabolism; galactose metabolism; sphingolipid metabolism; glutathione metabolism; amino acid and nucleotide metabolism; glycerolipid metabolism; starch and sucrose metabolism; and taurine metabolism. Additionally, there are other enriched pathways in the processing category, and subsequent categories like processing in signal transduction, cell differentiation and proliferation, repair proteins, and regulation of calcium concentration (e.g., FOXO signaling pathway, base excision repair, motor proteins, mTOR signaling pathway, and phosphatidylinositol signaling system; Table 2; Fig. 4C) that are essential for temperature responses.

**Table 2.** Significantly enriched Kyoto Encyclopedia of Genes and Genomes (KEGG) pathways for differentially expressed genes between the treatments in *Argiope bruennichi* (abru) spiderlings. # is the amount of enriched genes in the pathway.

| Pathway ID | Description | Category | # |
| --- | --- | --- | --- |
| abru00270 | Cysteine and methionine metabolism | Metabolism | 11 |
| abru00260 | Glycine, serine and threonine metabolism | Metabolism | 8 |
| abru00410 | beta-Alanine metabolism | Metabolism | 9 |
| abru00330 | Arginine and proline metabolism | Metabolism | 10 |
| abru00500 | Starch and sucrose metabolism | Metabolism | 12 |
| abru00380 | Tryptophan metabolism | Metabolism | 10 |
| abru01240 | Biosynthesis of cofactors | Metabolism | 26 |
| abru00591 | Linoleic acid metabolism | Metabolism | 8 |
| abru00561 | Glycerolipid metabolism | Metabolism | 14 |
| abru01200 | Carbon metabolism | Metabolism | 22 |
| abru00480 | Glutathione metabolism | Metabolism | 15 |
| abru00564 | Glycerophospholipid metabolism | Metabolism | 19 |
| abru00590 | Arachidonic acid metabolism | Metabolism | 13 |
| abru00760 | Nicotinate and nicotinamide metabolism | Metabolism | 7 |
| abru00190 | Oxidative phosphorylation | Metabolism | 17 |
| abru04142 | Lysosome | Cellular Processes | 35 |
| abru04814 | Motor proteins | Cellular Processes | 33 |
| abru04144 | Endocytosis | Cellular Processes | 34 |
| abru04145 | Phagosome | Cellular Processes | 24 |
| abru04137 | Mitophagy – animal | Cellular Processes | 22 |
| abru04146 | Peroxisome | Cellular Processes | 18 |
| abru04512 | ECM-receptor interaction | Signaling | 12 |
| abru03018 | RNA degradation | Processing | 17 |
| abru03410 | Base excision repair | Processing | 9 |
| abru03013 | Nucleocytoplasmic transport | Processing | 20 |
| abru04141 | Protein processing in endoplasmic reticulum | Processing | 38 |
| abru03040 | Spliceosome | Processing | 32 |
| abru04120 | Ubiquitin mediated proteolysis | Processing | 27 |
| abru04068 | FoxO signaling pathway | Processing | 21 |
| abru04150 | mTOR signaling pathway | Processing | 24 |

## Discussion

We assessed the vulnerability and degree of phenotypic plasticity of a cold-adapted *Argiope bruennichi* population from the northern expanding edge of the species’ distribution by exposing spiderlings in the egg sacs to three winter treatments. Despite strong temperature differences between treatments, the winter regimes did not affect spiderlings’ survival probability and energy stores (body lipids), indicating remarkable temperature tolerance. This aligns with previous findings where northern spiders had equal or higher survival proportions in warm winter temperatures (Wolz et al., 2020; Sheffer et al., 2024, preprint) to their own overwintering temperature. However, amounts of omega-3 polyunsaturated fatty acids, such as α-linolenic acid (ALA, C18:3n-3) were significantly reduced after three months of exposure in the warmer temperature compared to the cold, suggesting increased metabolic activity that can result in decreased lipid reserves when the spiderlings emerge from the egg sacs in the spring (Irwin & Lee, 2003). Subsequently, these physiological responses were supported by observed strong differences in gene expression between the treatments, specifically in response to the moderate and warm winter regimes. This might suggest a more targeted response to the cold regime, probably due to their cold adaptation at the northern edge.

Our study incorporated temporal information on survival proportion and fat content over winter, which showed that the temporal shifts have a more pronounced effect than treatment differences. Specifically, we observed a decrease in fat reserves and survival proportion from one to three months of treatment exposure, indicating the consumption of energy stores over time and subsequently resulting in lower survival. Other studies have reported a similar decline in survival proportions from mid to late winter, where post-winter mortality was associated with mass loss during the course of winter (Knapp & Řeřicha, 2020; Nielsen et al., 2022) that can severely impact post-winter fitness (Irwin and Lee, 2003). Fat reserves during winter are therefore biologically important, as lipids are the primary source of energy in overwintering insects (Hahn & Denlinger, 2011, Sinclair & Marshall, 2018). As ectotherms with temperature-dependent physiological responses (Sinclair, 2015), spiders are likely subject to similar trends, relying on lipids as their primary energy source. Both warmer temperatures and seasonal progression strongly influence overwinter energy use. For example, Potts and coauthors (2020) found a steady decrease of total lipids during the winter in the wolf spider *Schizocosa stridulans*. While we did not encounter differences in the overall lipid content at the end of the three months, we did find differences in fatty acid composition between treatments.

The roles of the fatty acid profiles in the metabolism during the overwintering period are well known in insects (Sinclair & Marshall, 2018.) However, in spiders, research in this area is still incipient. Our results revealed stable patterns of saturated and monounsaturated fatty acids as well as omega-6 polyunsaturated fatty acids (PUFAs) across the treatments. Nevertheless, we observed a step decline in omega-3 PUFA in spiders exposed to warm winters suggesting increased metabolic activity. In contrast, the spiderlings subjected to the cold treatment retained the omega-3 PUFAs, as these fatty acids may ensure membrane fluidity at low body temperatures (Sinclair & Marshall, 2018).

Our results can be explained by the nature of the fatty acids groups in different environments. Omega-3 PUFAs are highly unsaturated, possessing multiple double bonds that create kinks in their chains that prevents tight packing. This property is crucial for maintaining membrane fluidity in cold environments. However, in warm environments, the need for such high unsaturation decreases because higher temperatures naturally increase membrane fluidity. In contrast, omega-6 PUFAs have fewer double bonds and are less unsaturated than omega-3 PUFAs. This makes them more stable across a range of temperatures, as they do not disrupt membrane structure as much (Somero et al., 2017). Therefore, animals may maintain similar levels of omega-6 PUFAs regardless of environmental temperature.

In all three winter treatments, we found significantly more unsaturated fatty acids than saturated fatty acids. This aligns with the findings of Sinclair & Marshall (2018), who proposed that cold-adapted overwintering insects use this strategy to lengthen torpor and reduce metabolism. The consistent ratio of unsaturated/saturated fatty acids across treatments suggests a similar impact on fatty acid composition. This is expected, as ectotherms commonly adjust their lipid composition to ambient temperature to counteract harmful thermal effects on lipid fluidity (van Dooremalen et al., 2011). Our results also confirmed that spiders differ from other terrestrial arthropods like insects by having higher proportions of C20 polyunsaturated fatty acids (Uscian & Stanley-Samuelson, 1994; Stanley & Kim, 2020). Furthermore, the C20 PUFAs make up substantial proportions of the fatty acids associated with phospholipids, that play a structural role in membranes and are precursors of eicosanoids that play an important role as signaling and regulatory molecules (Wrońska et al., 2023). Because the spiderlings do not feed during the overwintering period, they must biosynthesize their eicosanoids from precursors that are provided by the mother in the egg yolk. Possibly, that is why they are constant across the winter regimes, as they are needed after the emergence from the egg sac, and do not play a role in the overwintering period.

To investigate the molecular mechanisms underlying the observed physiological responses, we analyzed gene expression patterns across the different temperature treatments. The majority of the differentially expressed genes (DEGs) found in our study displayed upregulation rather than downregulation. We also observed a higher overall transcriptomic response to increased temperature conditions, consistent with responses to temperature stress. Similar studies have shown that temperature stress on a newly cold-adapted population, typically triggers an active genetic response rather than the silencing of baseline gene expression (Lancaster et al., 2016; Tonione et al., 2020).

Additionally, we observed a substantial increase in the transcriptomic response to the moderate treatment, which could indicate adaptations or responses specific to the treatment, possibly involving different regulatory pathways. The increased expression in the moderate treatment can be explained by the nature of the chosen temperature treatments. The winter treatments emulated daily fluctuating temperatures that remained constant throughout the overwintering period (three months). Such daily fluctuations are typical in temperate climates but can severely compromise metabolic and physiological activity in arthropods (Xu et al., 2017). The temperature variations were extreme in both the cold and warm treatments, with a 20°C difference every day (Fig. 1), while the moderate treatment had only a daily 10°C difference. Williams et al. (2012) described for overwintering ectotherms how high daily thermal variability lowers the thermal sensitivity of increased metabolic rate due to having daily warmer temperatures thresholds. Daily temperature fluctuation can change the optimum and the critical maximum temperatures as well as the thermal reaction norms of a species (Paaijmans et al., 2013).

The moderate treatment elicited more diverse and less specific gene responses to thermal stress likely because the spiderlings were more temperature-sensitive. In contrast, the cold and warm treatments elicited more specific responses to cold and heat stress, respectively, showing the spiderlings in both treatments had lower thermal sensitivity. Production or breakdown of heat shock proteins or cryoprotectants, can further aggravate the influence of temperature variation, especially towards the extremes of the thermal regimes (Paaijmans et al., 2013).

Another aspect that contributes to the high degree of temperature tolerance in cold-adapted *Argiope bruennichi* spiderlings in the present study, might be explained by their range expansion history: Organisms were shown to remember their ancestral environments through phenotypic plasticity (Ho et al., 2020). *A. bruennichi,* from the northern edge of the distribution, originally from the Mediterranean, exposed to warmer winter temperatures might exhibit plastic reactions in gene expression that relate to their ancestral environment were winter temperatures are higher.

To gain insights into the functions of the differentially expressed genes, we performed a gene ontology (GO) analysis. The DEGs were most enriched in biological processes, specifically metabolic processes. The main energy sources for the overwintering spiderlings are primarily lipids, followed by glucose and some amino acids. Genes that affect energy production and other aspects of organismal function are expected to be upregulated when subjected to novel environmental conditions (Vasquez et al., 2019). This is indeed the case here, as the most enriched GO terms are from the warm and moderate temperatures, reflected in the in the number of genes associated with the GO terms (Table 1). The number of genes involved in the category is indicative of their biological importance to the response. Thus, the increment in functional enrichment for the moderate treatment indicates a rather unspecific response to temperature stress compared to the other treatments. In the moderate treatment, we found general stress responses, such as phosphorus metabolic processes, homeostatic processes, and negative regulation of biological processes in the Biological processes (BP) category. Other stress responses in the Molecular Function (MF) category that involved a high number of genes were Transferase activity and adenyl nucleotide binding ATP binding as seen in similar studies (L.-J. Chen et al., 2023).

The Kyoto Encyclopedia of Genes and Genomes (KEGG) analysis also revealed that metabolic pathways (including carbon, amino acid, nucleic acid, and fatty acid metabolism) were the most enriched and therefore further confirm that energy metabolism is critical for both cold and heat resistance (Sinclair, 2015; Mikucki & Lockwood, 2021). The visualization of the most significantly enriched biological pathways (Fig. 3c) provides insights into the predominant biological responses of *Argiope bruennichi* under different temperature conditions, offering a foundation for further functional characterization and investigation. The results also reaffirm the idea that the same set of genes can function differently depending on the exposure temperature, as evidenced by their significantly contrasting regulation. For instance, ribosomal expression is downregulated in the cold treatment (and upregulated in warm and moderate), which has been previously associated with diapause (Robich et al., 2007). The downregulation of the ribosomal expression in response to cold stress has also been found in in *Argiope bruennichi* from northern populations (Krehenwinkel et al., 2015), which could be another strategy to adjust protein production and slow down metabolic activity. Another example is the strong upregulation of sphingolipid metabolism in the warm and moderate treatments and its downregulation in the cold treatment. Sphingolipids (SLs) are major components of the plasma membrane and are important in the regulation of the thermal stress responses. SLs can structurally reinforce the membrane or/and send signals intracellularly to control numerous cellular processes (Fabri et al., 2020).

In addition to the previously discussed pathways, we also observed significant changes in Arachidonic acid metabolism. We found that Arachidonic acid metabolism is upregulated in warmer temperatures and downregulated in the cold. Arachidonic acid (ARA) and its derivatives play a crucial role in linking nutrient metabolism to immunity and inflammation. It is known that heat stress upregulates arachidonic acid to trigger autophagy (Hasan et al., 2019). Another reason why insects may have lower quantities of long-chain (C20) PUFAs, including ARA (Stanley & Kim, 2020), is that these PUFAs react with reactive oxygen species (ROS) generated by high oxidative catabolism and undergo peroxidation, which in turn leads to various forms of cellular damage (Lalouette et al., 2011).

While our study focused on the immediate responses to temperature variation during winter, it is important to consider the evolutionary adaptations that may occur during range expansions. Interestingly, we found a significant overlap of DEGs between cold and warm treatments. This suggests that the response to these temperature extremes involves a specific set of genes with distinct responses (upregulation vs downregulation), potentially highlighting unique biological pathways or stress responses activated by these temperature regimes. Some studies have documented an increase in cold tolerance during expansions to colder regions, but no change in heat tolerance and even a downregulation of genes involved in the heat shock responses. This is likely due to relaxed selection on heat tolerance when coping with colder environments (Kolbe et al., 2004; Krehenwinkel & Tautz, 2013; Lancaster et al., 2016). Furthermore, Lancaster et al. (2016) reported that several genes upregulated in response to heat stress in core populations became upregulated in response to cold stress in edge populations. This is because heat shock proteins can also prevent protein denaturation at cold temperatures. Heat shock proteins assist with folding and establishing proper protein conformation under stressful conditions, including cold conditions.

This study provides valuable insights into the molecular mechanisms underlying thermal tolerance and adaptation in *Argiope bruennichi* at its expanding edge. Our comparative transcriptome analysis of *A. bruennichi* spiderlings in different winter regimes provides data for the discovery of these mechanisms and uncovers the putative genetic adaptation of the species at the northern edge to the cold. The overall increased expression observed in the warm and moderate treatments compared to the cold demonstrates that *Argiope bruennichi* at the expanding edge is indeed adapted to colder environments. Their ability to alter the expression of metabolic pathways in different temperatures may contribute to their potential to survive rapidly changing environmental conditions.

*Argiope bruennichi* spiderlings in the cold-adapted population at the northern edge of the distribution are robust to different temperature winters. The ability to survive winter is highly critical to the distribution of species, so a thorough understanding of winter biology is crucial for predicting future responses to environmental change (Overgaard & MacMillan, 2017). We can affirm that the wasp spider has developed local adaptation and adaptive plasticity at the northern edge in response to current climatic conditions. Due to the lower dispersal probability of *A. bruennichi* at the expanding edge (Wolz et al., 2020), they must cope with increasing temperatures at northern latitudes or with even temporary colder conditions that can be caused by reduction of the snow cover. Our study shows that the spiderlings regulate gene expression and fatty acid composition according to the temperature regime. The observed changes in fatty acid profiles and gene expression, despite consistent physiological performance, suggest that *A. bruennichi* may employ compensatory mechanisms to maintain essential functions under varying thermal conditions. Nevertheless, these changes may ultimately affect their survival probability after emergence from the egg sac in the spring, particularly following warmer winters. Our study, highlights the importance of considering both physiological and molecular responses when assessing the vulnerability of species to climate change.

## Authors contributions

G.U. and M.K. conceived the study. G.U. sampled the population. M.W and M.K set up and conducted the temperature experiments. M.W, M.K and COM contributed to the performance data. AK, LJ, CJ and COM contributed to the transcriptomic data. AW and COM conducted the fatty acid analysis. COM analyzed all the data. COM wrote the original draft. COM, GU, and AW wrote and edit it the manuscript. All authors contributed critically to the drafts and read and approved the final manuscript.

## Acknowledgments

We want to thank Heidi Land and Christin Park, for their technical assistance. Lisa Hagenau, Jenny Melo, Zaide Ortiz and Laura Fuchs on their help with the analysis of transcriptomic technicalities. Andreas Fischer for his help with the GLMMs. The members of the General and Systematic Zoology group at the University of Greifswald provided support and helpful comments on the results.

