## Supplementary material for "Winter temperature effects in a cold-adapted northern population of a range-expanding spider: survival, energy stores, and differential gene expression": Tab 1

**Table S1.** Fatty acids amounts (mean ± SE; ng per individual) of the spiderlings from the three treatments. Only the fatty acids accounting for > 1% in at least one sample are shown. Σ SFAs is the sum of all saturated fatty acids. Σ MUFAs is the sum of all monounsaturated fatty acids. Σ ω-3 PUFAs is the sum of all poly-unsaturated omega-3 fatty acids, and Σ ω-6 PUFAs is the sum of all polyunsaturated omega-6 fatty acids. U/S is the ratio of unsaturated over saturated fatty acids. In red are the significant values of ANOVA and post Tukey HSD tests.

| Fatty acid | Treatments | | | $F_{2,21}$ | ANOVA, p | Tukey HSD | | |
|---|---|---|---|---|---|---|---|---|
| | **Cold(c)** | **Moderate(m)** | **Warm(w)** | | | **m -c** | **w- c** | **w- m** |
| C16:0 | 856.8 ± 360.7 | 758.2 ± 320.9 | 709.5 ± 241.1 | 0.465 | 0.635 | 0.804 | 0.618 | 0.948 |
| C18:0 | 1728 ± 510 | 1512.5± 400.8 | 1443.8 ± 529.8 | 0.752 | 0.484 | 0.652 | 0.480 | 0.957 |
| C20:0 | 80.6 ± 17.2 | 75.6 ± 22.0 | 71.46 ± 24.90 | 0.358 | 0.703 | 0.890 | 0.679 | 0.922 |
| C22:0 | 243.3 ± 418.2 | 93.9 ± 56 | 141.2 ± 19.7 | 0.741 | 0.489 | 0.471 | 0.698 | 0.925 |
| C24:0 | 9.2 ± 17 | 14.7 ± 20.44 | 14.3 ± 19.7 | 0.209 | 0.813 | 0.831 | 0.855 | 0.999 |
| **Σ SFA** | 2918 ± 1073.7 | 2455 ± 739.9 | 2380 ± 846.2 | 0.843 | 0.445 | 0.566 | 0.467 | 0.984 |
| C16:1n-9 | 97.8 ± 63.0 | 92.5 ± 53.5 | 101.81 ± 51.2 | 0.055 | 0.947 | 0.981 | 0.989 | 0.942 |
| C16:1n-7 | 131.5 ± 150.4 | 102.7 ± 73.8 | 165.3 ± 102.9 | 0.611 | 0.552 | 0.868 | 0.824 | 0.522 |
| C18:1n12 | 3785.7± 2782.1 | 3579.3 ± 2096.8 | 3203.9 ± 1177.7 | 0.154 | 0.858 | 0.979 | 0.849 | 0.934 |
| C18:1n-9 | 690.6 ± 964.3 | 276.9 ± 200 | 419 ± 344.3 | 0.974 | 0.394 | 0.372 | 0.645 | 0.885 |
| C20:1n-9 | 19.1 ± 16.2 | 14.9 ± 14.7 | 23.6 ± 24.9 | 0.407 | 0.671 | 0.902 | 0.887 | 0.645 |
| C22:1n-9 | 92.1 ± 236.7 | 11.9 ± 17.9 | 25.5 ± 13.7 | 0.782 | 0.470 | 0.485 | 0.603 | 0.979 |
| C24:1 | 28.1± 22 | 27.7± 24.7 | 34.6 ± 11.8 | 0.291 | 0.751 | 0.100 | 0.799 | 0.779 |
| **Σ MUFA** | 5015.3 ± 2791.2 | 4256.2 ± 2363.5 | 4097.3 ± 1585.6 | 0.363 | 0.700 | 0.789 | 0.708 | 0.990 |
| C18:4 n-3 | 120.9 ± 104.9 | 77.2 ± 51.2 | 43.9 ± 17.6 | 2.564 | 0.101 | 0.419 | 0.084 | 0.600 |
| C18:3 n-3 | 917.2 ± 588 | 502 ± 373 | 170.3 ± 122.7 | 6.721 | **0.005** | 0.129 | **0.004** | 0.257 |
| C20:3 n-3 | 14.4 ± 10.3 | 8.2 ± 7.1 | 15.9 ± 16.3 | 0.955 | 0.401 | 0.562 | 0.960 | 0.406 |
| C20:5 n-3 | 368.1 ± 143.9 | 367.2 ± 138.2 | 351.5 ± 161.2 | 0.032 | 0.969 | 0.100 | 0.972 | 0.976 |

| | | | | | | | | |
|---|---|---|---|---|---|---|---|---|
| C20:4 n-3 | 128.6 ± 183.3 | 133.8 ± 137.5 | 53.1 ± 34 | 0.913 | 0.417 | 0.997 | 0.507 | 0.462 |
| C22: 5 n-3 | 27.9 ± 19.9 | 19 ± 16.2 | 31.9 ± 6.9 | 1.486 | 0.249 | 0.484 | 0.866 | 0.235 |
| C22:6 n-3 | 15.6 ± 21.9 | 4.9 ± 13.9 | 22.4 ± 22.2 | 1.585 | 0.228 | 0.533 | 0.778 | 0.206 |
| **Σ ω-3 PUFA** | 1592.8 ± 734.6 | 1112.4 ± 503.2 | 689.2 ± 313.6 | 5.504 | **0.012** | 0.206 | **0.008** | 0.288 |
| C18:2 n-6 | 2582.3 ± 1915.8 | 2350.8 ± 1524.3 | 3125.5 ± 1171.7 | 0.515 | 0.605 | 0.953 | 0.769 | 0.592 |
| C18:3 n-6 | 595.6 ± 943.7 | 519.8 ± 828.4 | 107.2 ± 95.2 | 1.045 | 0.369 | 0.976 | 0.388 | 0.504 |
| C20:2 n-6 | 22.6 ± 42.9 | 17.3 ± 22.1 | 20.1 ± 24.3 | 0.059 | 0.943 | 0.937 | 0.986 | 0.982 |
| C20:3 n-6 | 43.2 ± 57.5 | 19.8 ± 22.5 | 24.4 ± 22.5 | 0.851 | 0.441 | 0.449 | 0.589 | 0.969 |
| C20:4 n-6 | 225.9 ± 138.6 | 211.0 ± 94.4 | 342.8 ± 150.9 | 2.455 | 0.11 | 0.971 | 0.195 | 0.131 |
| C22: 2 n-6 | 107.8 ± 80.4 | 83.7 ± 22.8 | 85.3 ± 28.4 | 0.558 | 0.581 | 0.617 | 0.659 | 0.998 |
| **Σ ω-6 PUFA** | 3577.5 ± 1406.2 | 3202.5 ± 802.4 | 3705.4 ± 1288.1 | 0.383 | 0.686 | 0.807 | 0.975 | 0.682 |
| **U/S** | 3.5 | 3.5 | 3.6 | 0.208 | 0.814 | 0.928 | 0.959 | 0.799 |
| **TOTAL FA** | **13103.6± 4541.2** | **11026.1 ± 3803.8** | **10872.3± 3596.3** | **0.776** | **0.473** | **0.561** | **0.515** | **0.997** |

**Table S2**. Fisher's Exact Test for differentially gene expressed (DGEs) data.

| | Odds ratio | p-value |
|---|---|---|
| cold vs moderate | 1.11538 | 0.6121 |
| cold vs warm | 0.65566 | 0.0068 |
| warm vs moderate | 0.03801 | 0.00001 |

**Table S3**. Loadings of the first two differential expressed genes (DEGs) principal components of the for each RNAseq sample.

| Sample | PC1 | PC2 | Treatment |
|---|---|---|---|
| Cold_S1 | -3.623337 | 10.1330979 | cold |
| Cold_S2 | -2.966298 | 3.9336629 | cold |
| Cold_S3 | -4.937319 | 1.4266976 | cold |
| Cold_S4 | -4.421512 | 4.8106767 | cold |
| Moderate_S5 | -9.422837 | 0.9821716 | moderate |
| Moderate_S6 | -17.518834 | -10.1148940 | moderate |
| Moderate_S7 | -12.287558 | -1.6018582 | moderate |
| Moderate_S8 | 9.835132 | 1.3030997 | moderate |
| Warm_S9 | 11.173751 | -3.2870693 | warm |
| Warm_10 | 12.066212 | -2.9077292 | warm |
| Warm_11 | 11.220850 | -3.0356624 | warm |
| Warm_12 | 10.881751 | -1.6421932 | warm |

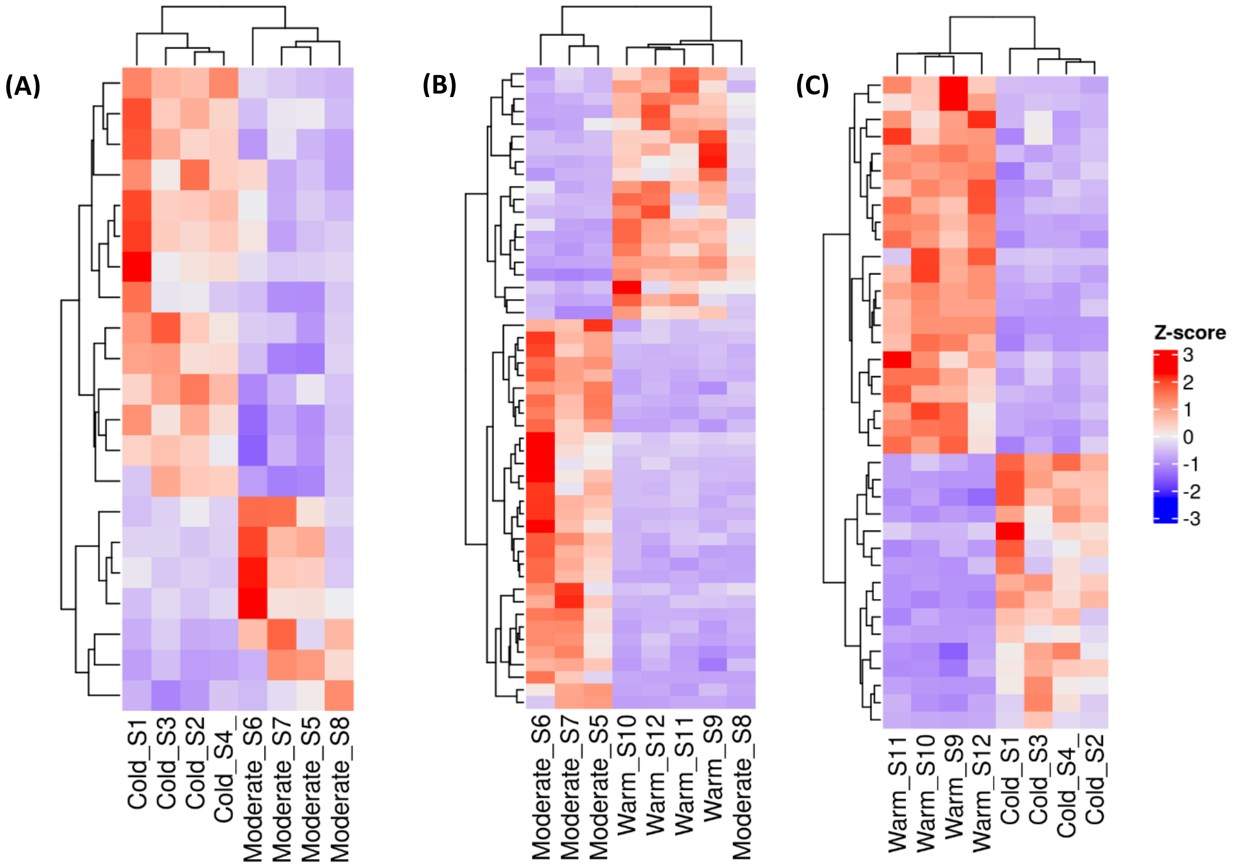


**Figure S1. Heatmaps of normalized gene expression values of significant DEGs**. **(A)** moderate vs cold, **(B)** warm vs moderate and **(C)** cold vs warm. The Z-score represents the normalized gene expression value. Red indicates upregulated expression and blue indicates downregulation expression. Columns correspond to samples and the rows are genes.

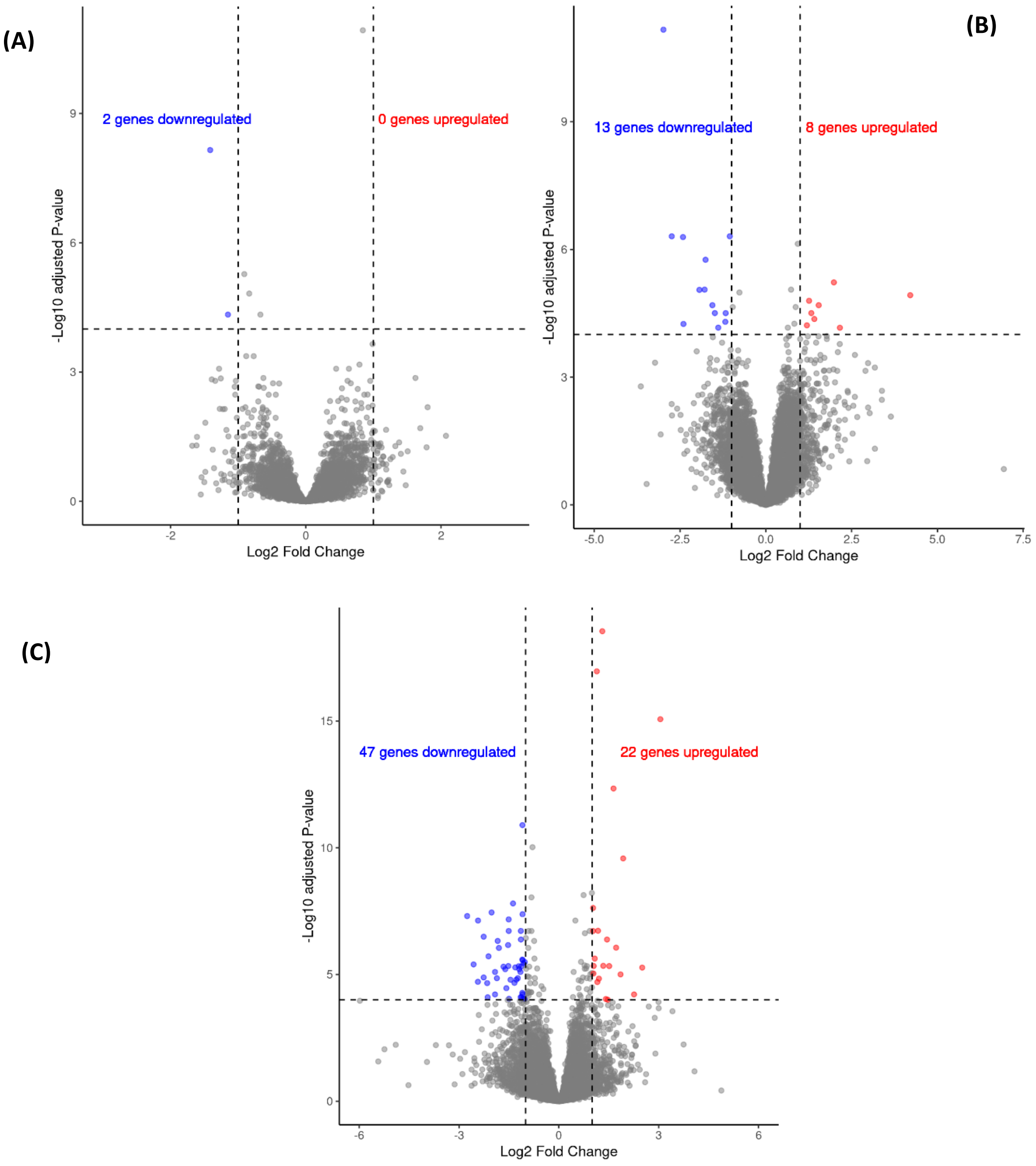


**Figure S2. Volcano plots of significantly DEGs between (A)** moderate vs cold, **(B)** warm vs moderate and **(C)** cold vs warm. No significant genes are represented in grey. Highly significant (p-adj) upregulated genes in red and downregulated in blue.
